# Heightened condition dependent expression of structural colouration in the faces, but not wings, of male and female flies

**DOI:** 10.1101/2021.06.09.447681

**Authors:** Thomas E. White, Amy Locke, Tanya Latty

## Abstract

Structurally coloured sexual signals are a conspicuous and widespread class of ornament used in mate choice, though the extent to which they encode information on the quality of their bearers is not fully resolved. Theory predicts that signalling traits under strong sexual selection as ‘honest’ indicators should evolve to be more developmentally integrated and exaggerated than nonsexual traits, thereby leading to heightened condition dependence. Here we test this prediction through examination of the sexually dimorphic faces and wings of the cursorial fly *Lispe cana*. Males and females possess structural UV-white and golden faces, respectively, and males present their faces and wings to females during close-range, ground-based courtship displays, thereby creating the opportunity for mutual inspection. Across a field-collected sample of individuals, we found that the appearance of the faces of both sexes scaled positively with individual condition, though along separate axes. Males in better condition expressed brighter faces as modelled according to conspecific flies, whereas condition scaled with facial saturation in females. We found no such relationships for their wing interference pattern nor abdomens, with the latter included as a nonsexual control. Our results suggest that the structurally coloured faces, but not the iridescent wings, of male and female *Lispe cana* are reliable guides to individual quality and support the broader potential for structural colours as honest signals. They also highlight the potential for mutual mate choice in this system, while arguing for one of several alternate signalling roles for wing interferences patterns among the myriad taxa which bear them.

## Introduction

Colour patterns present a striking dimension of phenotypic variation, and nowhere is this better showcased than in the context of sexual communication. The variable ornaments of male guppies (Endler 1991; Houde 1987), iridescent signals of butterflies (Kemp 2008a; White et al. 2015), and exaggerated badges of hummingbirds (Greenwalt et al. 1960) are exemplars and have each served as models for examining the role of sexual selection in driving the evolution of conspicuous visual signals. A central hypothesis is that such signals are selectively favoured as honest guides to the genetic and/or phenotypic quality of potential mates, with empirical tests primarily guided by costly-signalling and index models (reviewed in Weaver et al. 2017). Costly-signalling models, such as the Zahavian handicap, predict costs to signal production or maintenance which are differentially borne among signallers (Grafen 1990; Zahavi 1975). Among-individual differences in the ability to acquire resources underlie differences in their ultimate allocation, with only the ‘best‘ individuals able to produce and bear the most brilliant signals. Indices, by contrast, describe how signal production is unfakably tied to the function of internal processes (Maynard-Smith 2003). The energy, resources, and/or time required for signal production are not costly unto themselves under such an explanation, and honesty is instead maintained by direct links to core physiological processes. The expected outcome of both processes, which stands as the key test of theory, is that signals should exhibit heightened-condition dependent expression as compared to traits under weaker sexual selection (Cotton 2004a).

Almost all colour signals in nature are the product of absorption by pigments or scattering by nanostructures (Johnsen 2012). Empirical tests of honesty-based models have chiefly focused on the former, with carotenoid-based ornaments receiving particular attention (reviewed in Blount & Mcgraw 2008; Svensson & Wong 2011). As pigments that cannot be synthesised *de novo* carotenoids must be acquired through diet (Blount & Mcgraw 2008). This environmental dependence creates opportunity for selection to favour links between resource acquisition and allocation and, ultimately, signal expression. The red plumage of the house finch *Haemorhous mexicanus* offers a well characterised example, with recent work revealing how the yellow-to-red bioconversion of dietary carotenoids prior to deposition links individual condition (via mitochondrial efficiency) to the quality of visual displays (Hill et al. 2019), which are used to inform mate choice (Hill 1994).

Structural colours, by contrast, arise from by an interaction between light and nanostructures that vary in refractive index, and are capable of degrees of brilliance and spectral richness otherwise unattainable through pigments alone (Vukusic & Sambles 2003). Despite their widespread use as conspicuous sexual ornaments, the case for honesty in structurally colour signals is less well developed. There are three broad arguments regarding such potential. For one, if the construction and/or maintenance of nanostructures is materially demanding, then this may create a trade-off against other core needs (Keyser & Hill 1999; Zahavi 1975). Such demands will then be differentially met among individuals of varying quality, as consistent with a handicap-based explanation (Zahavi 1975).

A second argument rests on the precision with which nanostructures must be arranged for optimal signal expression, and hence their sensitivity to perturbation during development (Ghiradella & Butler 2009). If individuals vary in the stability of environmental conditions (e.g. thermal or nutritional) experienced during development, either incidentally or as the result of active choice, then the resulting signals may act as an index of phenotypic and/or genetic quality (Ghiradella & Butler 2009; Shawkey et al. 2003).

Finally, the accumulating evidence of self-assembly during structural colour development (e.g. Prum et al. 2009; Maia et al. 2011) has underlain arguments against any expectation of condition-dependence (Prum 2009), assuming the absence of active and expensive cellular processes. This latter assumption appears inconsistent with recent work, however (Rubenstein et al. 2021), and the broader weight of evidence supports the scaling of structural colour expression with measures of mate ‘quality’ (reviewed in White 2020), as well as mate choice based on such variation (e.g. Kemp 2008; Kodrick-Brown & Johnson 2002). Though valuable, this body of work remains heavily taxonomically biased toward birds, and more often than not lacks the nonsexual control necessary for tests of heightened condition-dependent expression (Cotton 2004a), thereby limiting the strength of and generality of inferences which may be drawn.

Flies rank among the most diverse animal orders and showcase striking adaptations to support their visually rich lives (Lunau 2012; Marshall 2012). A relatively poor colour sense across the Diptera has historically implied a limited capacity or need for colour-mediated communication (Troje 1993), but work in select species continues to document the use of visual ornaments and dynamic displays in the service of mate choice (eg. Butterworth et al. 2019; Butterworth et al. 2021; Zimmer et al. 2003). To that end, recent attention has centred on ‘wing interference patterns’ (WIPs) as visual displays and the targets of sexual selection (Hawkes et al 2019; Katayama et al. 2014). These conspicuous patterns adorn the semi-transparent wings of many insects, including flies, and are a product of thin-film interference at the air/chitin interface(s) of wing membranes (Shevtsova et al. 2011). Our understanding of their possible role as signals is nascent, but evidence for their active presentation during courtship (e.g Frantsevich & Gorb 2006; White et al. 2019), heritability (Hawkes et al 2019), and evolutionary lability in response to sexual selection (Katayama et al. 2014; Hawkes et al. 2019) is consistent with their use as signals, with the encoding of information on mate quality being one plausible, but untested, function.

*Lispe cana* is a cursorial species of muscid fly endemic to supralittoral habitats spanning the entire Eastern coast of Australia (Pont 2019). They possess sexually dimorphic, structurally coloured faces and WIPs, the latter of which exhibit limited-view iridescence. These conspicuous patterns are actively presented during distinctive courtship displays in which males pursue females, before engaging in a ritualised ground-based ‘dance’ at close range (Frantsevich & Gorb 2006; White et al. 2019). The clear potential for both male and female assessment during courtship offers a promising context for testing the potential for honesty in structurally coloured ornaments, which formed the motivating aim of our study. As discussed below, such colours in holometabolous (completely metamorphic) insects are constructed and fixed during ontogeny from the pool of resources gathered during the larval stage (Hunt 2004; Rowe 1996). This means that a field sample of adult phenotypes offers a population-level statement of condition and signal expression that effectively integrates all underlying environmental and genetic influences on each. The key prediction for our field-based study, then, was for heightened condition dependence in the structurally coloured faces and wings of both male and female *Lispe cana*, under the hypothesis that such ornaments function as indicators of mate quality.

## Methods

### Field sampling

We collected 47 female and 57 male *Lispe cana* from the supralittoral zone of Toowoon bay, New South Wales, Australia (33.3626° S, 151.4975° E). We humanely euthanised all collected individuals by chill-coma *in situ* using a refrigerated esky, before transporting them to a laboratory at The University of Sydney, Camperdown, Australia for processing, as described below. We preserved all specimens in a refrigerator at a maximum of 2° to prevent the degradation of structures and/or pigments, and we took all measurements within three weeks of capture.

### Assessment of condition and colour traits

In holometabolous insects the adult phenotype—including colour signals and body size— is constructed from the resources acquired during the larval stage and fixed at eclosion. Since the quality and quantity of larval resources define the ‘quality’ of the resulting phenotype—as closely indicated by adult body size—this total pool of resources can be considered equivalent to individual condition (Hunt 2004; Rowe 1996). We therefore used adult body size, indicated by thorax length, as a surrogate measure of condition, which is also typical of past work in flies (e.g. Cotton et al. 2004b; David et al. 2000). We used scaled digital images of collected flies to measure the distance between the anterior prothorax and posterior metathorax in imageJ (Rueden et al. 2017).

To quantify signal expression, we measured the reflectance of three body regions across both male and female flies: their structurally coloured faces and wings, and their black, melanic abdomens. Abdomens were included as a trait whose visual appearance is assumed to not be under sexual selection (given it is unviewable during courtship), which is an important control for testing the heightened condition dependence predicted by indicator models (Cotton 2004; White 2020). Prior to measurement we non-destructively separated the heads and wings of flies from the thorax and mounted each region on a ca. 90 × 90 mm square of matte-black card. We used an OceanInsight JAZ spectrometer with pulsed PX-2 Xenon light source, coupled with a 400 μm bifurcated probe to both send and collect light which we oriented at 45° relative to sampling surfaces. We aligned faces and wings with their dorsal and anterior edges nearest the probe, respectively. This setup gave a ca. 4 mm sampling spot size which encompassed the frons and vertex of faces and spanned the entirety of the central wing region between the terminus of the subcostal vein on the anterior margin and the anterior cubital vein on the posterior margin. We used a Spectralon WS-1 as a diffuse white standard and recalibrated against it between each measurement.

To estimate the chromaticity and luminance of signals as relevant to potential mates, we used a slightly amended form of the dipteran visual model of Troje (1993). We drew on the visual phenotype of the muscid fly *Musca domestica* as the nearest available analogue to *Lispe cana*, and assumed the involvement of R7p, R8p, R7y, and R8y photoreceptors in chromatic processing, and R1-6 in achromatic processing (Hardie 1986; Troje 1993). For chromatic contrasts we estimated receptor quantum catches as the integrated product of stimulus reflectance, an ideal (i.e. flat across the 300-700 nm range) illuminant, and each receptor’s sensitivity function, before calculating the difference in relative stimulation between R7y-R8y and R7p-R8p receptors. These two putative opponent channels define the location of a given stimulus in two-dimensional dipteran colourspace, from which we took the Euclidean distances between a stimulus and the achromatic centre as our measure of saturation (or chroma). We estimated luminance as the absolute stimulation of R1-6 receptors, following the estimation of quantum catches as above. We conducted all spectral processing and visual modelling in R (v 4.1.0; R Core Team) using the packages ‘lightr’ (v1.1; Gruson et al. 2019) and ‘pavo’ (v 2.7.0; Maia et al. 2019).

### Statistical analysis

We used generalised linear models fit by restricted maximum-likelihood to test the prediction of heightened condition dependence across six signalling traits: the chromaticity and luminance of faces, wings, and abdomens. Each trait served as a response, and we specified the interaction between sex and condition (body size) as predictors in all models, with the latter representing the key test of condition-dependence. We specified a Gaussian error distribution with identity link function for all models, and visually confirmed the assumptions of additivity and residual normality. We also standardised all parameter estimates by centring predictors to have a mean of zero and dividing by their standard deviations for ease of comparison and interpretation (Gelman 2008). All statistical analyses were carried out in R (v 4.1.0; R Core team 2021).

### Data availability

All underlying data and code will be persistently archived upon acceptance.

## Results

Facial colouration in *Lispe cana* is strongly sexually dichromatic (Fig. 1a) and condition-dependent (Fig. 2a, b). The dichromatism stems from males exhibiting considerably brighter faces than females by virtue of their broadband UV-white reflectance. By contrast, the golden-yellow appearance of female faces is characterised by a sigmoidal-type reflectance with an inflection at ca. 520 nm, which underlies their heightened chromaticity as compared to the achromatic faces of males (Table 1). We saw little evidence for dichromatism in wing interference patterns, though this may in part be a consequence of our measuring at whole-wing scales. The weakly multi-modal reflectance profiles of wings (Fig. 1b) are a product of the contributions of individual wing panels which vary in thickness and, hence, chromaticity and brightness. That is, the mosaic of conspicuously chromatic panels are relatively achromatic, and sexually monomorphic, at whole-wing scales (but see discussion for further detail).

**Table 1:** Summary descriptors of the visual characteristics of male and female faces and wing interference patterns in *Lispe cana*. Chroma and luminance were estimated according to a colourspace model considering the visual system of conspecific flies, and abdominal measures are included as a nonsexual control in our tests for heightened condition-dependent expression in signalling traits. Values represent means ± standard errors.

| Parameter | Males (n = 57) |  | Females (n = 47) |  |
| --- | --- | --- | --- | --- |
|  | Mean | Range | Mean | Range |
| Facial chroma | 0.015 ± 0.001 | 0.001 - 0.047 | 0.308 ± 0.008 | 0.207 - 0.550 |
| Facial luminance | 1.260 ± 0.045 | 0.551 - 1.801 | 0.427 ± 0.021 | 0.074 - 0.689 |
| Wing chroma | 0.045 ± 0.005 | 0.005 - 0.188 | 0.041 ± 0.005 | 0.005 - 0.019 |
| Wing luminance | 0.226 ± 0.022 | 0.075 - 0.398 | 0.226 ± 0.016 | 0.098 - 0.499 |
| Abdominal chroma | 0.012 ± 0.002 | 0.002 - 0.032 | 0.003 ± 0.001 | 0.053 - 0.031 |
| Abdominal luminance | 0.022 ± 0.001 | 0.008 - 0.057 | 0.023 ± 0.002 | 0.008 - 0.042 |

**Figure 1:**
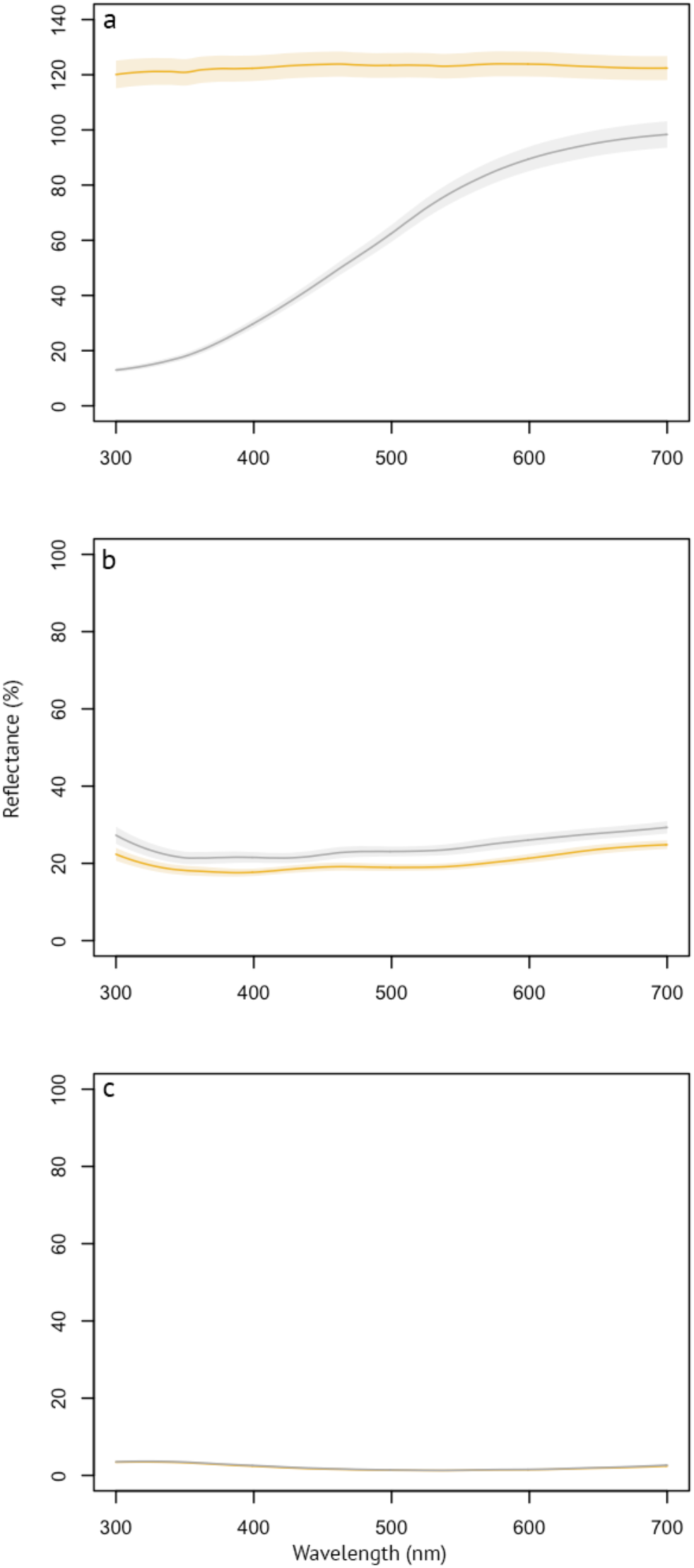
Reflectance spectra (mean ± s.e.) of the (a) faces, (b) wing interference patterns, and (c) abdomens of male (grey) and female (gold) *Lispe cana*. Note that males and females are near-completely overlain in (c).

**Figure 2:**
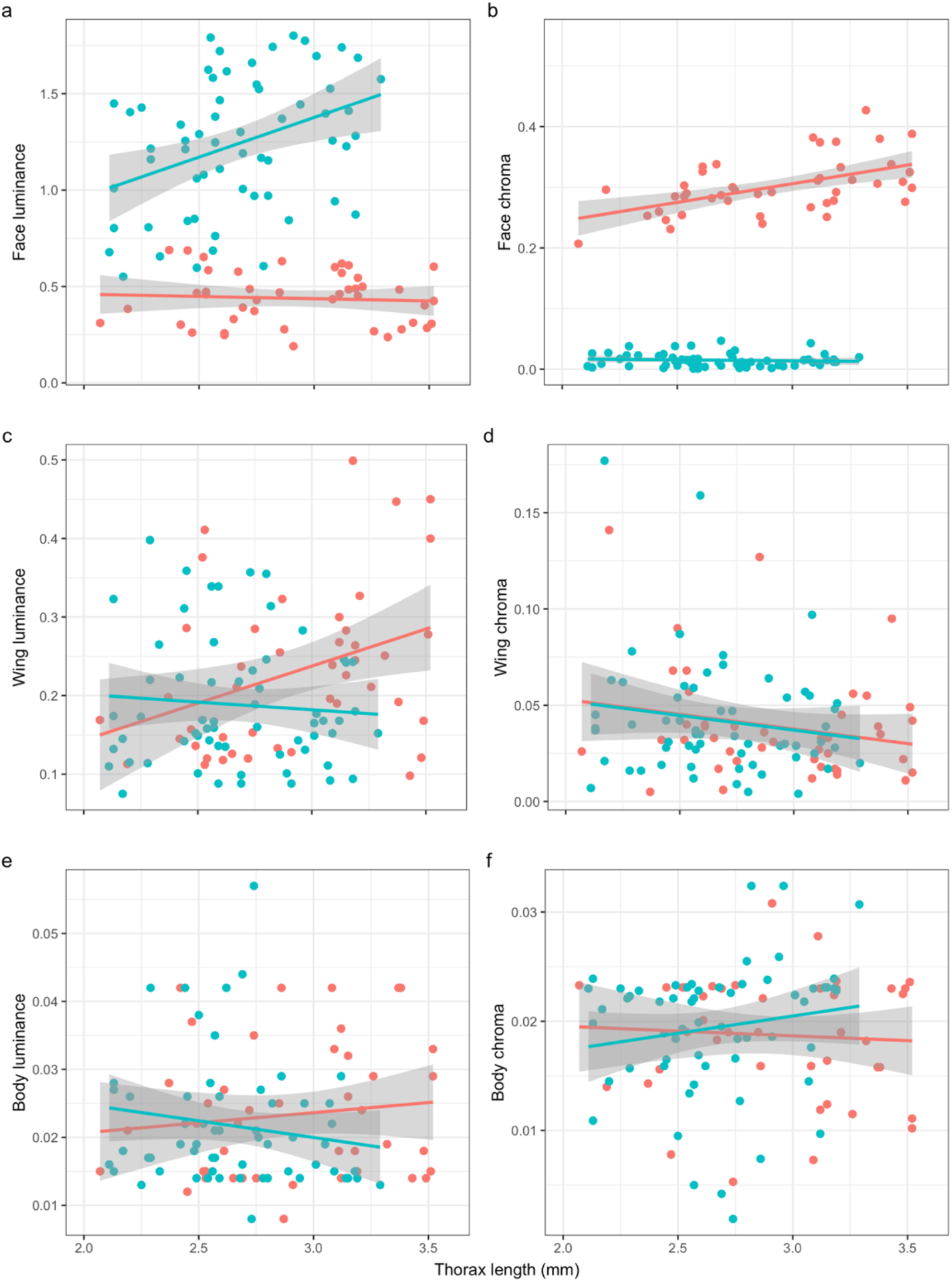
Raw data and generalised linear model fits describing the relationship between colour signal expression and individual condition in *Lispe cana* (n = 47 females, 57 males). Shown are estimates of the luminance and chromaticity of the (a, b) faces, (c, d) wings, and (e, f) abdomens against thorax length as a measure of condition, for both male (green) and female (red) flies.

We identified significant condition dependence in the faces of both males and females as indicated by the sex by size interaction. It manifested along separate axes in each sex (Table 2). The faces of larger males are more luminant across the 300-700 nm range (Fig. 2a), whereas the faces of larger females are characterised by increased chromaticity (Fig. 2b). The reciprocal did not hold, hence the interaction, with no apparent relationship between male condition and facial chromaticity, nor female condition and facial luminance. The WIPs of both sexes bore no relationship to body condition along any dimension, nor did their abdomens as our nonsexual control (Fig. 2c-f).

**Table 2:** Standardised parameter estimates and test statistics from six generalised linear models testing for the condition-dependent expression of structural colouration in the faces and wings of the fly *Lispe cana*. All models included sex (male/female), condition (via thorax length), and their interaction as predictors, specified with a Gaussian error distribution and identity link function. Bolded estimates are statistically significant at α = 0.05.

| Response | Sex | Condition | Interaction |
| --- | --- | --- | --- |
| Facial chroma | $\beta = -0.28 \pm 0.01, t = -40.18, p < 0.001$ | $\beta = 0.02 \pm 0.01, t = 3.28, p = 0.001$ | $\beta = -0.06 \pm 0.01, t = -4.47, p < 0.001$ |
| Facial luminance | $\beta = 0.86 \pm 0.06, t = 15.47, p < 0.001$ | $\beta = 0.17 \pm 0.06, t = 2.88, p = 0.005$ | $\beta = 0.40 \pm 0.11, t = 3.46, p < 0.001$ |
| Wing chroma | $\beta = 0.00 \pm 0.01, t = 0.24, p = 0.810$ | $\beta = -0.01 \pm 0.01, t = -0.892, p = 0.374$ | $\beta = -0.00 \pm 0.02, t = -0.217, p = 0.829$ |
| Wing luminance | $\beta = -0.03 \pm 0.02, t = -1.70, p = 0.09$ | $\beta = 0.01 \pm 0.02, t = 0.34, p = 0.732$ | $\beta = -0.05 \pm 0.04, t = -1.44, p = 0.154$ |
| Abdominal chroma | $\beta = -0.01 \pm 0.01, t = 0.837, p = 0.40$ | $\beta = 0.01 \pm 0.02, t = 1.07, p = 0.287$ | $\beta = 0.00 \pm 0.01, t = 1.08, p = 0.281$ |
| Abdominal luminance | $\beta = -0.01 \pm 0.01, t = -1.04, p = 0.299$ | $\beta = 0.00 \pm 0.02, t = -1.11, p = 0.269$ | $\beta = -0.01 \pm 0.03, t = -0.90, p = 0.370$ |

## Discussion

Structurally coloured ornaments are often-extravagant products of sexual selection, though evidence for their role as ‘honest’ indicators of mate quality is heterogeneous (White 2020). Here we examined the key prediction of condition dependence in the structurally coloured faces and wing interference patterns of the cursorial fly *Lispe cana*. We found evidence for the moderate to strong scaling of facial signal expression with body size—a proxy measure of condition—in both sexes, albeit along distinct axes. Males in better condition were brighter, while females were more chromatic, and no such relationship was apparent for wing interference patterns in either sex. Comparison against a nonsexual control supported the contention of heightened condition dependence among these putative signalling traits. Though observational, our results affirm the potential for structurally coloured ornaments to serve as informative signals of mate quality, while identifying opportunities for mutual mate choice on complex multi-dimensional ornaments.

The sexual differences we identified in facial colouration and the axes of condition-dependence are underlain by differences in physical mechanisms. The bright UV-white faces of males are the product of incoherent scattering by disordered nanostructures, as is true of non-fluorescent white colours in nature in general (Johnsen 2012; Vukusic et al. 2003; Wiersma 2013). In *L. cana*, the scattering elements are densely packed scales which are modified into flat, elongated bristles (ca. 60 × 6 um) during development (unpublished data; but see Frantsverch & Gorb 2006 for details in closely related species). Although the nanostructural basis of variation within sexes remains to be described, theory (Johnsen 2012) and empirical work (Frantsverch & Gorb 2006) supports the primacy of bristle density as a predicted determinant of the among-male variation in facial brightness here identified (Fig. 2a), with further possible contributions from bristle geometry and any internal structuring. That is, the sheer number of scattering elements will chiefly distinguish higher from lower quality individuals, and hence the availability and quality of material gathered during the larval stage are a plausible limiting resource. Analogous dynamics are well described in other holometabolous insects, such as the pierid butterfly *Eurema hecabe*. Males display an iridescent ultraviolet wing patch, the brightness of which is driven, in part, by the density of reflective elements adorning individual wing scales (White et al. 2012). The arrangement of these elements is susceptible to perturbation through manipulations of the quality of larval foodplant. Male signal brightness therefore offers a window to juvenile foraging success and developmental environments, which females use to inform their choice of mate (Kemp 2008a; Kemp 2008b).

Female facial colouration in *L. cana* shares the same fundamental bristle-based architecture as males, though their golden hue is imparted by the addition of pigments studded across the facial surface. At a proximate level, the condition-dependent variation in saturation we identified (Fig. b) will be driven by the quantity of underlying pigments and the density of reflective structures acting in concert. More pigments mean a greater fraction of shorter-wavelength incident light will be absorbed, leading to increased spectral purity (Johnsen 2012). Similarly, greater broadband scattering by bristles will increase the relative reflection of longer versus shorter wavelength light, and so will also increase saturation, albeit to a lesser degree.

A mechanistic understanding of the links between female condition and signal expression awaits identification of the pigments in use in *L. cana*, though carotenoids and pterins are likely candidates. The former is dietarily acquired and the latter synthesised *de novo*, and each have been implicated as signals of quality (Walker et al. 2019; Weiss et al. 2011). Irrespective of the proximate cause, however, the potential content of such signals is clear in light of the well-recognised scaling of female body size and fecundity in insects (Honek 1993). Male choosiness is expected to be favoured where substantial variance in female quality exists, as suggested here (Fig. 2), and when the costs to mate searching and assessment are low (as in the flies’ high-density foreshore habits) but mating itself are high (Bonduriansky 2001). These conditions appear well met in *L. cana*, and males stand to benefit from discriminating among females on the basis of facial saturation, though whether and to what extent they do so remains to be seen.

Unlike faces, we found no evidence for condition dependence among the wing interference patterns of either sex. This is unsurprising among females given their wing patterns are never actively displayed and are unlikely to be incidentally seen by conspecifics. The absence of an effect among males however, for which a signalling role for WIPs is likely, suggests two possibilities. One is our measurements did not capture signal variation at the functionally relevant spatial, spectral, or temporal scale. Wing interference patterns are a mosaic of panels which are delineated by wing venation. The colours of each element are chiefly defined by the thickness and spacing of the air/cuticle multilayer, as well as any surface structuring such as ridges or bumps (Shevtsova et al. 2011). At whole-wing scales, such as those measured here, the wings of *L. cana* appear to be only weakly chromatic as the contributions of these individual panels average out across the visible spectrum (Fig. 1b). Although this represents the experience of most viewers under most conditions, male *L. cana* actively present their wings at a distance of only ca. 5 mm during courtship. The visual acuity of *Lispe* is unknown, though data from related species (e.g. 5.0 cycles per degree in the muscid *Musca domestica*; Land 1972) suggests the possibility that individual wing panels may be spatially resolvable at these signaller/receiver distances common to courtship. In which case the appearance of particular wing regions and/or their spatial arrangement may bear salient information on male quality, the signal of which would be masked at whole-wing scales such as those considered here.

By a similar token, males’ striking wing patterns are never viewed in stasis. Males rapidly ‘flutter’ their wings during their ritualised courtship dances and move in rapid lateral semi-circles around females who are constantly reorienting in response (White et al. 2020). This presentation behaviour suggests a role for the temporal structure of signals as a channel of information. Modifications to the corrugation of wings and/or the arrangement of surface structures (such as microtrichia; Shevtsova et al. 2011) to enhance or suppress limited-view iridescence, for example, may be similarly indicative of resource limitation or broader developmental stress, as discussed above. Yet such variation would only be apparent to us through the measurement of wing signal angularity (which was beyond the scope of the present work), and to conspecific viewers through the active presentation of wings during courtship. There is morphological and behavioural evidence in insects (Kemp et al. 2006; White et al. 2015) and birds (Stavenga et al. 2011) which indirectly supports the possibility, though it remains an intriguing working hypothesis for future study.

The second broad possibility is that wing interference patterns do not function as indicators and instead fulfill one of many other potential roles during signalling. Numerous insects, including flies, are attracted to flashing stimuli (Eichorn 2017; Magnus 1958), with work in butterflies showing this preference can increase linearly up unto the limits of temporal resolution (Magnus 1958). A male’s rapidly flickering wings may therefore serve to capture and hold a female’s attention during courtship, or bias subsequent gaze directions toward their luminant and centrally located faces. A second, related, possibility is that male WIPs serve as amplifiers of the true foci of female choice (Hasson 1991; Byers et al. 2010). Their faces are an obvious candidate, though the environmentally contingent nature of WIPs means that the behavioural performance of males during courtship could also be readily assessed. The limited-view structure of interference patterns displayed on semi-transparent wings means that optimal colour expression (or any colour expression at all) is only achievable via presentation against suitably dark backgrounds and under sufficiently specular lighting. Male *L. cana* can and do exert some active control over each by biasing the microhabitats in which they display (White & Latty 2020; White et al. 2020). Thus, if a male’s ability to select suitable microhabitats varies with some facet of individual quality, then the appearance of WIPs would render such information apparent to female viewers. This would be a novel form of visual signal amplification enabled by direct ties to display environments, though evidence for the broader phenomenon is well established (reviewed in Byers et al. 2010).

Our results support a growing, albeit heterogeneous, body of evidence supporting the potential for honesty among structurally coloured ornaments (e.g. Griggio et al. 2010; Kemp 2008b; McGraw et al. 2002). This was true of both sexes in our focal system which suggests the potential for mutual mate choice, and also extends the male-biased focus in this (White 2020) and related (e.g. Ah-King et al. 2014) areas of research. That we found no evidence for heightened condition dependence in WIPs narrows the scope of explanations for the adaptive evolution of these widespread ornaments (Shevtsova et al. 2011). A complete understanding, however, awaits a richer appreciation of the spectral, spatial, and temporal complexity of WIPs, and colour-based signals more generally. Exciting theoretical work continues to advance these aims at several levels (e.g. Stoddard & Osorio 2019; van den Berg et al. 2020), and tractable systems such as *Lispe* sp. hold excellent promise for empirical progress.

## Acknowledgments

TEW thanks Elizabeth Mulvenna and Cormac White for their endless support.

